# Inhibition of Mitochondrial Fission by Drp-1 Blockade by Short-Term Leptin and Mdvi-1 Treatment Improves White Adipose Tissue Abnormalities in Obesity and Diabetes

**DOI:** 10.1101/2021.07.23.453523

**Authors:** P Finocchietto, H Perez, G Blanco, V Miksztowicz, C Marotte, C Morales, J Peralta, G Berg, C Poderoso, JJ Poderoso, MC Carreras

**Affiliations:** Laboratorio de Metabolismo del Oxígeno INIGEM-UBA-CONICET, Buenos Aires, Argentina; Departamento de Medicina, Facultad de Medicina, Universidad de Buenos Aires, Buenos Aires, Argentina; Laboratorio de Inmunotoxicología (LaITo), IDEHU-CONICET, Universidad de Buenos Aires; Facultad de Medicina, Pontificia Universidad Católica Argentina (UCA), Instituto de Investigaciones Biomédicas (UCA-CONICET), Laboratorio de Patología Cardiovascular Experimental e Hipertensión Arterial, Buenos Aires, Argentina; Laboratorio de Lípidos y Aterosclerosis, Departamento de Bioquímica Clínica, Facultad de Farmacia y Bioquímica, Instituto de Fisiopatología y Bioquímica Clínica (INFIBIOC), Universidad de Buenos Aires, Buenos Aires, Argentina; Departamento de Patología, Facultad de Medicina. Instituto de Fisiopatología Cardiovascular, Universidad de Buenos Aires; Departamento de Bioquímica Humana, Facultad de Medicina, Universidad de Buenos Aires

**Keywords:** white adipose tissue, mitochondria, biogenesis, dynamics, Drp1, mdivi-1

## Abstract

**Background:** Obesity and type 2 diabetes are chronic diseases characterized by insulin resistance, mitochondrial dysfunction and morphology abnormalities.

**Objective:** Herein, we investigated if dysregulation of mitochondrial dynamics and biogenesis is involved in an animal model of obesity and diabetes.

**Methods:** The effect of short-term leptin and mdivi-1 –a selective inhibitor of Drp-1 fission-protein– treatment on mitochondrial dynamics and biogenesis was evaluated in epididymal white adipose tissue (WAT) from male *ob/ob* mice.

**Results:** An increase in Drp-1 protein levels and a decrease in Mfn2 and OPA-1 protein expression were observed with enhanced and sustained mitochondrial fragmentation in *ob/ob* mice compared to *wt C57BL/6* animals (p<0.05). The content of mitochondrial DNA and mRNA expression of PGC-1α –both parameters of mitochondrial biogenesis– were reduced in *ob/ob* mice (p<0.05). Leptin and mdivi-1 treatment significantly increased mitochondrial biogenesis, improved fusion-to-fission balance and attenuated mitochondrial dysfunction, thus inducing white-to-beige adipocyte transdifferentiation. Measurements of glucose and lipid oxidation in adipocytes revealed that both leptin and mdivi-1 increase substrates oxidation while *in vivo* determination of blood glucose concentration showed decreased levels by 50% in *ob/ob* mice, almost to the *wt* level.

**Conclusions:** Pharmacological targeting of Drp-1 fission protein may be a potential novel therapeutic tool for obesity and type 2 diabetes.

## INTRODUCTION

Obesity and type 2 diabetes are chronic diseases that typically coexist and have grown in prevalence to become a global health problem. Both of them share a pathophysiological axis of insulin resistance (IR), oxidative stress, mitochondrial dysfunction and chronic inflammation. Obesity is characterized by an abnormal increase in white adipose tissue (WAT) mass depicting an imbalance between energy intake and consumption that leads to energy overload (1,2).

In mammals, there are three types of adipose tissue: white, brown and beige, with distinct functions and morphology, different protein expression patterns, and dissimilar developmental origins (3,4). While the function of WAT is to store energy in the form of lipid droplets that can be released to fuel other tissues, brown adipose tissue (BAT) has thermogenic properties for maintaining body temperature (5-7). Browning is the process by which some adipocytes within WAT depots acquire properties of brown adipocytes (an increase in mitochondrial content and oxidative metabolism) showing an intermediate phenotype between white and brown adipocytes (“beige” or “brite”) (8).

Leptin, the satiety and anti-obesity hormone, is an adipokine released from white fat tissue that acts on the hypothalamus to induce satiety and reduce food intake, and on the peripheral tissues to control metabolism and to increase the metabolic rate (9).

*Ob/ob* mice suffer a mutation that determines a premature stop codon in the *Lep* gene (leptin) and constitute a well-known model of obesity and type 2 diabetes characterized by hyperglycemia, hyperinsulinemia, high food intake, mitochondrial dysfunction, and oxidative/nitrosative stress affecting mitochondrial complex I activity in WAT, liver and skeletal muscle (10-11).

Mitochondria are dynamic organelles essential for cellular energy survival and a major site of ROS production. These organelles, whose morphology has a remarkable plasticity according to the energetic cell needs, undergo constant mitochondrial fusion and fission, and are replaced every 2–4 weeks in different tissues through mitophagy. All of these processes are necessary for mitochondrial quality control and clearance (12,13). Energetic homeostasis is sustained through the balance among mitochondrial biogenesis, dynamics, ultrastructure, function and degradation in a healthy mitochondrial population (14). While mitochondrial fusion is mediated by the mitofusin proteins (Mfn) 1 and 2 in the outer membrane and by OPA-1 (optic atrophy-1) in the inner membrane, mitochondrial fission requires the translocation of Drp-1, a member of the large GTPases dynamic family, from the cytosol to the mitochondrial outer membrane. When the cytosolic protein Drp-1 is activated, it translocates to the outer membrane of the mitochondrion where it multimerizes generating a ring-like structure that constricts and divides this organelle. Post-translational modifications of the protein contribute to the regulation of mitochondrial fission mainly by phosphorylation of serine residues that increase or decrease its GTPase activity (15). Drp-1 activation is rapidly regulated by the phosphorylation of serine 616 and dephosphorylation of serine 637, which are targeted by different phosphatases and kinases. While phosphorylation of Drp-1 on Ser-637 prevents its mitochondrial translocation, phosphorylation of Drp-1 on Ser-616 promotes mitochondrial fragmentation during mitosis (16).

Mdivi-1 is a quinazolinone that selectively inhibits Drp-1 over other dynamin family members and prevents mitochondrial fission by pharmacological inhibition. This drug inhibits Drp-1 self-assembly into rings and its association with mitochondria (17-18).

Mitochondrial biogenesis is the process through which new mitochondria are generated, driven by the transcriptional activators NRF-1 and 2 and by PGC-1 alpha. It is activated by various signaling pathways, including nitric oxide (NO)/cyclic GMP, Akt and AMPK, among others (19). The AMP-activated protein kinase (AMPK) is a cellular energy sensor that when activated by phosphorylation leads to upregulation of PGC-1α, UCP-1, and mitochondrial biogenesis (20).

Alterations of mitochondrial dynamics and function have been implicated in neurodegeneration, aging, sepsis, cardiovascular disease, obesity and type 2 diabetes (21-24). In this context, the twofold aim of the present study is to investigate if dysregulation of mitochondrial dynamics and biogenesis are involved in a relevant animal model of obesity and diabetes, and if Drp-1 can be pharmacologically targeted as a potential novel therapeutic tool for obesity and diabetes.

## MATERIALS AND METHODS

### Animals experimentation

Male *ob/ob* (± 70 g) and *wt C57BL/6* (± 25 g) 5-month-old mice were purchased from Jackson Labs (USA). Animal experiments were performed in accordance with the Principles of Laboratory Animal Care. The animal experiments were approved by the local Scientific and Technology Ethics Committee at the University of Buenos Aires (UBA). All efforts were made to minimize animal suffering and to reduce the number of animals used. Mice were maintained under controlled temperature (21° C +/-2° C), humidity (50–60%), and air-flow conditions, with a fixed 12-h light/dark cycle. Until the experiment onset, all animals were fed with a standard mice laboratory chow and had free access to food and water to standardize their nutritional status. Animals were sacrificed by cervical dislocation at the end of the treatment with leptin or mdivi-1.

### Drug administration

*Wt C57BL/6* and *ob/ob* mice received short-term (3 days) intraperitoneal injection of pyrogen-free dimethyl-sulfoxide, or recombinant leptin (1 mg/kg/d dissolved in dimethyl-sulfoxide) (L 3772 Sigma Chemical Co.), or mdivi-1 (50 mg/kg/d dissolved in dimethyl-sulfoxide) (Mdivi-1 MO199 Sigma Chemical Co). In all mice, body weight and food intake were measured (11,17).

### Blood glucose determination

On the day of sacrifice, blood samples were collected using cardiac puncture. Blood glucose concentration was determined by Accu-check performance glucometer (Roche Lab). Animals were fasted for 10h before any procedure.

### White adipose tissue extraction

Mice were sacrificed by cervical dislocation and epididymal white adipose tissue (6-10 g) was immediately extracted and homogenized in sucrose buffer 20% (TRIS 10 mM, EDTA 0.1mM, sucrose 20%, 2% protease inhibitor cocktail) (Sigma Aldrich, St. Louis, MO, USA). The homogenate was centrifuged at 800*g* for 10 min at 4°C, frozen in liquid nitrogen and stored at 80°C until further analysis (11,24).

### Western Blot Analysis

Proteins (50 *μ*g) were separated using electrophoresis on 7.5-10% SDS-polyacrylamide gel, and transferred to a PVDF membrane (GE, Healthcare). Membranes were incubated with antibodies against 1: 1000 anti-Mfn2 (H-68): sc-50331, 1: 4000 anti-actin (I-19): sc-1616 obtained from Santa Cruz, CA or 1: 1000 anti-OPA-1: 612607 and 1: 1000 anti-Drp-1: 611113 obtained from BD Biosciences, 1:1000 anti-LC3 II (L7543 Sigma-Aldrich), 1: 1000 anti-phospho-Drp1 (Ser616): #3455, 1: 1000 anti-phospho-Drp1 (Ser637): #6319, 1:1000 anti-AMPKα: #2532, 1:1000 anti-phospho AMPKα (Thr172): #2531 obtained from Cell Signaling. After several washes, the membranes were incubated with appropriate horseradish peroxidase conjugated secondary antibodies. Detection of immunoreactive proteins was accomplished by enhanced chemiluminescence. Quantification of bands was performed by digital image analysis using Total Lab Analyzer software (Nonlinear Dynamics Ltd, Biodynamics, Argentina). Equal loading was controlled with the appropriate markers (11,24).

### Electron microscopy images

WAT was fixed in 4% paraformaldehyde, 2% glutaraldehyde, and 5% sucrose in PBS, followed by 2-h post-fixation in 1% osmium tetroxide, and then 1 h in uranyl acetate in 50% ethanol. Samples were washed with ethanol 50% and dehydrated with a graded series of ethanol, clarified with acetone and embedded in Vestopal. Grids were prepared and stained with uranyl acetate and lead citrate. Samples were observed at 100 kV with a Zeiss EM-109-T transmission electron microscope (Zeiss, Oberkochen, Germany) (11,24).

### Optic microscopy images and histological evaluation

The histological examination by light microscopy was performed in a blinded manner. Fixed epididymal WAT samples were dehydrated in ethanol, embedded in paraffin wax, and cut with a microtome Reichert (Austria). The resulting micro-sections were stained with hematoxylin and eosin reagent and periodic acid-schiff (PAS) stain for the determination of size and density of adipocytes and vascular density. In both cases the quantification was performed at high power field (HPF), 20 fields at 400x magnification for each animal using a computerized image analyzer (Image Pro Plus, Media Cybernetics Corp) (25).

### Isolation of mitochondria

Mitochondria were isolated from homogenized WAT by differential centrifugation. Mitochondrial pellets were stored in the presence of antiproteases and antiphosphatases as described (11,24).

### Mitochondrial complex I activity

The activity of Complex I (NADH: ubiquinone reductase) was determined spectrophotometrically following the reduction of 50 μM 2,3-dimethoxy-6-methyl-1,4-benzoquinone at 340 nm by 50 μg/ml mitochondrial proteins with a Hitachi U3000 spectrophotometer at 30°C; 200 μM NADH was used as electron donor. Reaction was carried out in the presence of 1 mM KCN and expressed as nmol of reduced benzoquinone/min.mg prot. Complex I activity was selectively inhibited by 10 µM rotenone (11,24).

### Primary fat cell isolation

WAT pads were weighed and sliced into 3 mm pieces with scissors and resuspended in 5 ml of KRH buffer (25 mM NaHCO_3_, 12 mM KH_2_PO_4_, 1.2 mM MgSO_4_, 4.8 KCl, 120 mM NaCl, 1.4 mM CaCl_2_, 5 mM glucose, 20 mM Hepes, 2% protease inhibitor cocktail (Sigma Aldrich, St. Louis, MO, USA), pH7.4, plus 2.5% BSA containing 0.5 mg/ml collagenase (Sigma Chemical Co., St Louis, MO, USA). The fat pads were digested for 45-60 minutes at 37°C in an orbital bath shaking at 100 rpm. Adipocytes released from the tissue were harvested by centrifugation (400g, 15 min). Cells in the upper phase of the centrifuge tube were collected, washed three times and resuspended in 0.15 M PBS. Adipocytes were isolated from animals without treatment and under the different treatments (11,24).

### Fluorescence microscopy images and analysis

Adipocytes from mice under treatment were stained with mitotracker green (50uM) (Invitrogen Carlsbad, CA) and observed under an Olympus BX-51 fluorescence microscope equipped with a digital Q-Color 3 Olympus camera. Digital images were processed with Image-Pro Plus 6.0 software (Media Cybernetics, Rockville, MD, USA) and Cell Profiler (Broad Institute, USA). A Cell Profiler pipeline was created for the segmentation of individual mitochondria and measurement of morphometric parameters. Several images for each treatment were processed with this pipeline to identify and measure more than 300 mitochondria. All morphometric measurements of mitochondria were exported to a standard database and were further analysed with Cell Profiler Analyst (Broad Institute, USA). A standard random forest classification algorithm was used to implement a machine learning approach to classify mitochondria intro three groups: round, elongated and branched. The training procedure was repeated until global classification accuracy achieved 90%, with 100% classification accuracy of the round mitochondria subgroup. Then, the whole set was scored for the amount of mitochondria belonging to each of the three groups (26-28).

### Adipocytes siRNA transfection

Adipocytes were transfected with siRNAsDrp-1 or empty-vector siRNA (Santa Cruz Biotechnology) 50nM using lipofectamine (Invitrogen Corp. California, USA) in Opti-MEM reduced serum medium and were incubated at 37°C in 5% CO_2_ for 10 hours, according to the protocol provided by the manufacturer (11). Silencing efficiency (60%) was evaluated by Drp-1 immunoblotting (Suppl. Fig.2). The sequence of siRNAs Drp-1-sc-45953 was designed on the structure of mice genome (11,24).

### Oxidative metabolism in isolated adipocytes

Isolated adipocytes were incubated with leptin (200 ng/ml), mdivi-1 (50 uM) or siRNA Drp-1 (50nM) for 10h, and with ^14^[C] palmitic acid, samples were distributed into tubes containing Whatman filter paper soaked in NaOH and 200 nCi/ml ^14^[C] palmitic acid. The tubes were sealed and incubated for 2 h; then, 10 N HCl was added to release ^14^[C] CO, which was detected by scintillation counting of the filter paper. To measure complete glucose oxidation to CO_2_ and H_2_O, samples were supplemented with D-glucose ^14^[C[U]]-[250 mCi/mmol]. Radioisotopes were from Perkin-Elmer Life and Analytical Sciences, Boston, MA, USA (11,24).

### RNA Extraction and Real-Time PCR (RT-qPCR)

Total RNA was insulated from the different tissues using TriZol reagent following the manufacturer’s instructions (Life Technologies, Inc.-BRL, Grand Island, NY). Any residual genomic DNA was removed by treating RNA with RQ1 Rnase-free DNase (Promega, Madison, WI, USA) at 37°C for 30 min, which was subsequently inactivated by incubation with 2 mM EGTA for 10 min at 65°C. First-strand cDNA synthesis was performed using oligo(dT)18 primer, Invitrogen SuperScript IV Reverse Transcriptase (RT), and RNase inhibitor (RNasin, Promega, Madison, WI, USA). PCR amplification and analysis were performed with ABI PRISM 7500 Sequence Detector System (PE Applied Biosystems, Foster City, CA).

SYBR Select Master Mix (Applied Biosystems, Carlsbad, CA, USA) was used for all reactions, following manufacturer’s instructions. Real-time PCR data were analyzed by calculating the 2^-ΔΔCt^ value (comparative Ct method) using GAPDH expression as housekeeping, performed in parallel as endogenous control (11,24). The primer sequences and the condition of each reaction are shown in Supplemental Table I. **mtDNA/nDNA ratio**. WAT was homogenized in lysis buffer (10 mM Tris-HCl pH 8, 1 mM EDTA, and 0.1% SDS). After adding proteinase K, lysates were incubated at 55°C for 3 h, vigorously vortexed and centrifuged (8,000 g for 15 min); the resting supernatant was vortexed with 1 ml protein precipitation solution (Gentra Puregene Kit) for 30 sec and placed on ice for 5 min. The resulting supernatant was mixed with 1 volume of isopropanol and centrifuged (12,000 g for 15 min at 4°C) to precipitate DNA. The DNA pellets were washed with 70% ethanol, air dried, and dissolved in Tris-EDTA buffer. For mtDNA/nDNA qPCRs, 40 ng of total genomic DNA were used. The mtDNA/nDNA ratio was calculated using the formula: 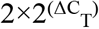, where ΔC_T_ is the difference of C_T_ values between nDNA ß2M and the mtDNA 16S rRNA primers as shown in (29).

## Statistical analysis

Data are presented as mean ± SEM according to the normal or skewed distribution. Significant differences between groups were accepted at p < 0.05. One-way ANOVA (multigroup comparisons) followed by Bonferroni’s multiple comparison test or Dunnett’s test were carried out with GraphPad Prism 5.01 (La Jolla, CA).

## RESULTS

### Short-term leptin and mdivi-1 treatment reverts mitochondrial morphological defects in WAT from *ob/ob* mice

Mitochondrial morphology defects were detected by TEM in WAT slices in *ob/ob* mice in comparison to *wt C57BL/6* and *ob/ob* treated with leptin or mdivi-1 mice. While *wt* mitochondria were predominantly tubular-shaped (80%), *ob/ob* mitochondria were aberrantly small and spherical in shape (85%) (Fig. 1 A and B). Leptin or mdivi-1 treatment resulted in a significant increase in tubular-shaped mitochondria (55% and 49%, respectively) (Fig. 1 C, D and E). As mitochondrial fragmentation and dysfunction lead to mitophagy, we analyzed WAT slices by TEM and observed mitochondria engulfed by double-membrane vacuoles with mitochondrial fragments indicating an active process of mitophagy in *ob/ob* mice (10 images per field). These alterations were reduced after treatment with leptin (5 images per fields) or mdivi-1 (3 images per field) (Fig. 1F). We also assessed the LC3 II protein levels, which increased in *ob/ob* mice and decreased with leptin or mdivi-1 treatment (Fig.1 G). No changes were observed in *wt C57BL/6*.

**Figure 1.**
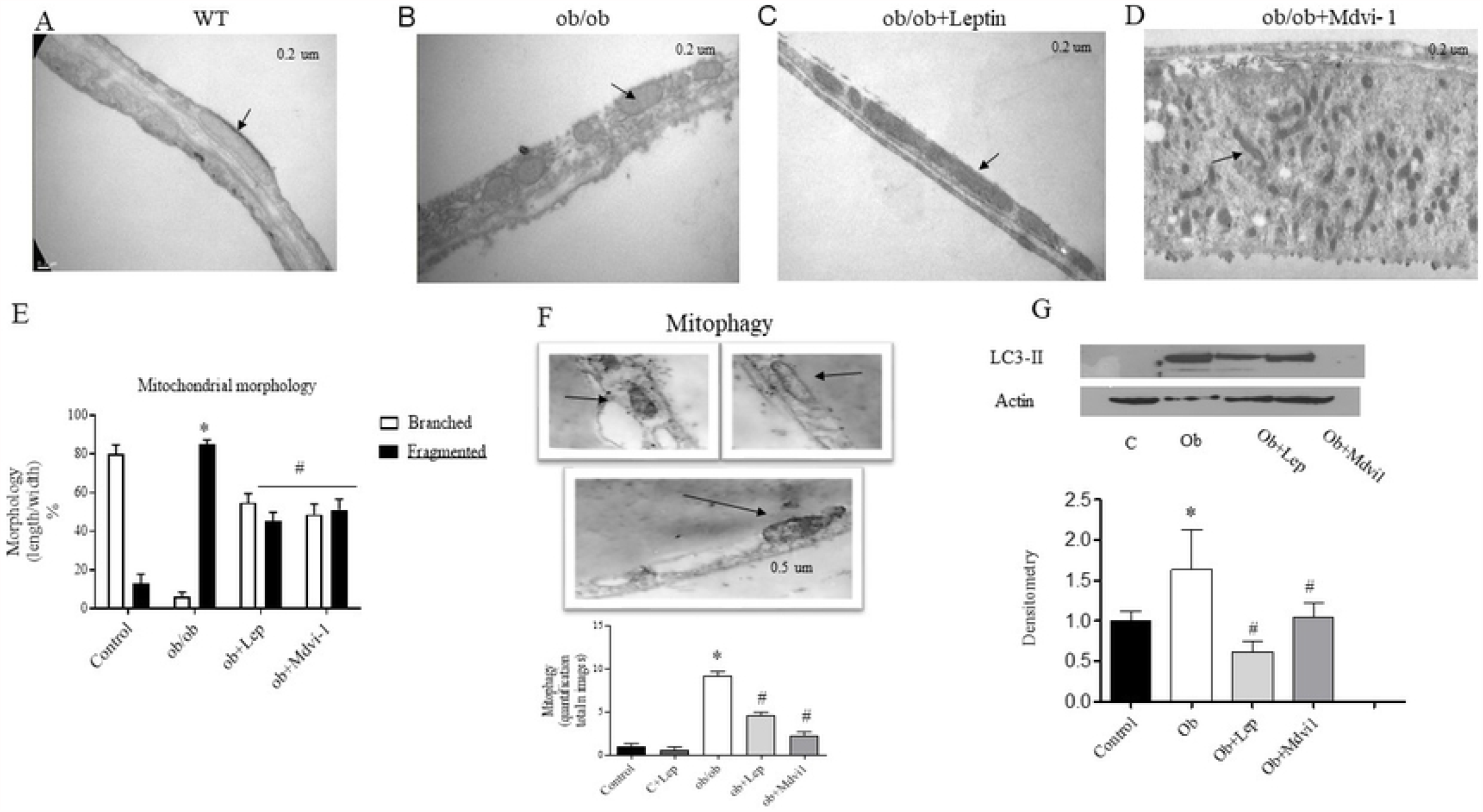
Effect of short-term leptin and mdvi-1 treatment on mitochondrial ultrastructural morphology. Representative electron microscope images of mitochondrial morphology were evaluated using electron microscopy of fixed white adipose tissue from *wt C57BL/6* (A) or *ob/ob* (B) (Magnification 3000-30.000 X) with leptin (C), or mdvi-1(D) treatments (bar represents = micrometer), (5 images per animal for 6-8 per group) (black arrows). Tubular and fragmented mitochondria were counted per arbitrary area. (E) The percentage distribution of tubular and fragmented adipocyte mitochondria was determined in a minimum of 8–10 random fields at 3.000-30.000 X magnification to ensure a representative area of analysis. Those mitochondria whose length were more than thrice its width were considered tubular while round mitochondria were considered fragmented. (F) Representative images of WAT *ob/ob* mice slices reveal the presence of mitophagy characterized by autophagic vacuoles with mitochondrial particles or mitophagosomes (black arrows) (total images of mitophagy in 10 images per animals). (G) Representative Western blots of LC3 II protein. Bars reflect the densitometry in arbitrary units (A.U.) in absolute amount of protein. Data are normalized to ² -actin for protein. Results are mean ± SEM, n= 5-7, ^*^p < 0.05 denotes different from respective *wt C57BL/6*. # p < 0.05 denotes different from respective *ob/ob, ob/ob* vs *ob/ob* + leptin or mdvi-1 treatment. n:5 per group.

### Short-term leptin and mdvi-1 treatment reverts mitochondrial morphological defects in isolated adipocytes from WAT

We investigated mitochondria morphological changes on isolated adipocytes from WAT *wt C57BL/6* or *ob/ob* mice treated with leptin or mdvi-1 intraperitoneally using a machine learning algorithm to allow automated classification of mitochondria shape into three groups: branched, round, or elongated detected by fluorescence microscopy images (Suppl. Fig. 1 A).”Branched” mitochondria denoted complex shapes including T-shapes, Y-shapes, P-shapes, O-shapes, among others, and have been associated with particular bioenergetic profiles (29). The results of mitochondrial classification into each of the three groups through machine learning is shown in Fig. 2 A and B. The measurement confirmed that *ob/ob* mice had a higher proportion of round mitochondria compared to *wt C57BL/6* mice (0.45±0.05 to 0.22±0.04), consistent with the TEM analysis in WAT slices. This proportion was partially restored in *ob/ob* mice treated with leptin (0.36±0.04), and completely restored or even decreased by mdvi-1 treatment (0.22±0.05).

**Figure 2.**
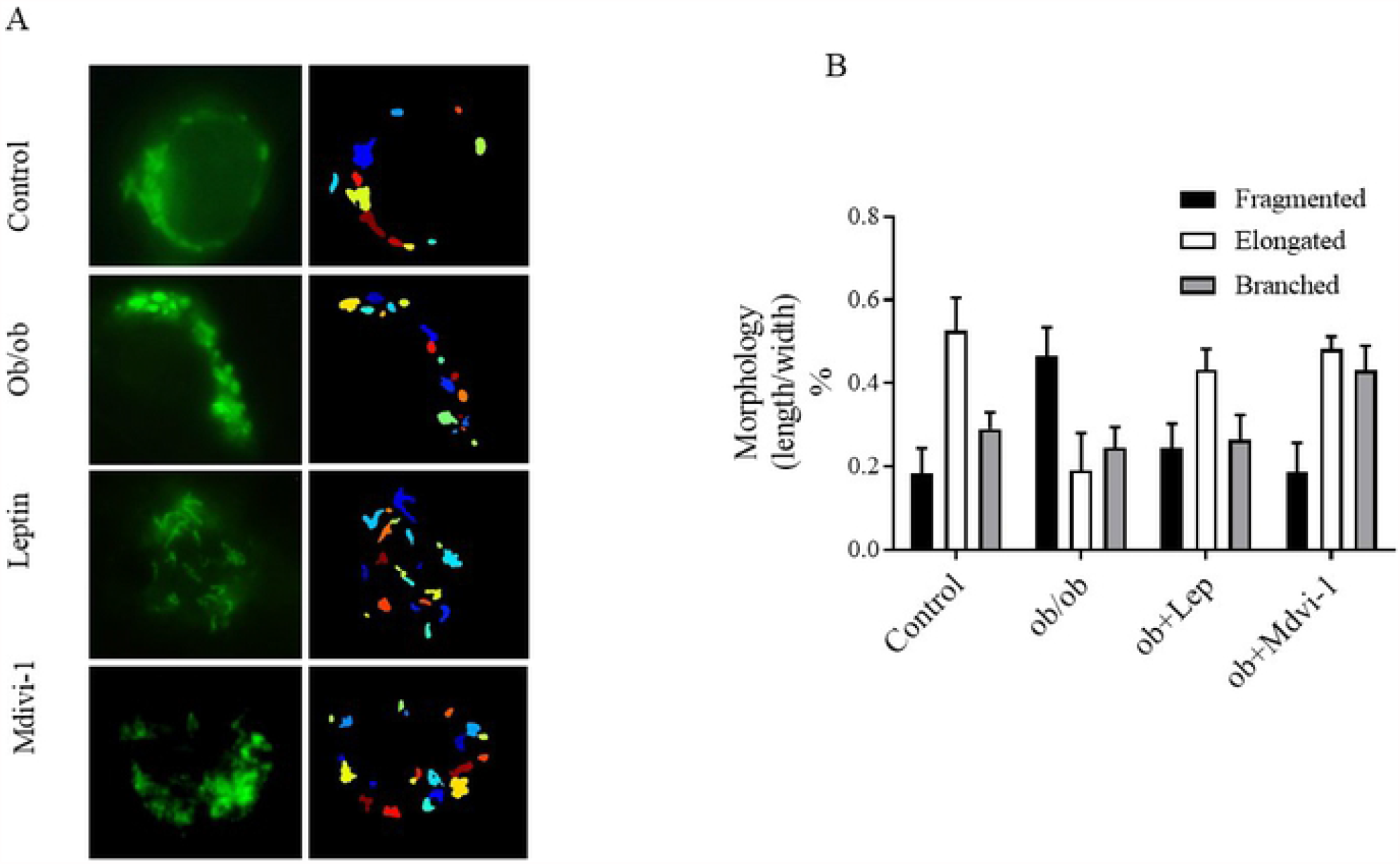
Effect of leptin and mdvi-1 on mitochondria shape-classes in adipocytes isolated from WAT. (A) Representative micrographs of adipocytes stained with mitotracker green and observed under fluorescence microscopy (digitally enlarged 1.000 x magnification), and their matching segmentation obtained with the Cell Profiler pipeline for extracting morphological measurements. Different mitochondrial colours mean mitochondrial morphology: blue and light blue: a branched, yellow: rounded, orange and red:elongated. (B) Proportions of the three defined mitochondria shape-classes among treatment groups with their corresponding 95% confidence intervals followed by supervised classification into three mitochondrial shape-classes (to further determine proportions of each class).

The proportion of elongated mitochondria was strongly reduced in *ob/ob* mice when compared against *wt C57BL/6* mice (0.19±0.04 and 0.47±0.06, respectively). Both leptin and mdvi-1 treatment increased the presence of elongated mitochondria (0.31±0.04 and 0.31±0.06, respectively). In contrast, *wt C57BL/6, ob/ob* and leptin-treated groups showed a similar proportion of branched mitochondria (0.31±0.06, 0.36±0.05 and 0.32±0.06, respectively), while mdvi-1-treated mice showed an increased proportion of branched mitochondria (0.46±0.05) (Fig. 2B and Suppl. Fig. 1A). No changes were found in *wt C57BL/6* under any treatment.

### Short-term leptin treatment modulates the mitochondria fission-to-fusion ratio in WAT from *ob/ob* mice

In order to examine whether leptin can modulate the mitochondria fission in leptin-deficient *ob/ob* mice with mitochondrial dynamics impaired, we analyzed the expression of the mitochondrial fission (Drp-1) and fusion (Mfn2 and OPA1) proteins in WAT with and without leptin treatment. Drp-1 protein level is increased by ∼50% in the WAT of *ob/ob* mice compared to *wild type* animals (Fig. 3A). Drp1 activity is regulated by the opposing effects of phosphorylation at two key serine residues: phosphorylation of Ser 616 increases Drp-1 activity whereas it is decreased by phosphorylation of Ser 637 (30). We measured the phosphorylation of both residues and found an increase in p-Ser 616-Drp-1 (active form) whereas p-Ser 637-Drp-1 (inactive form) decreased in *ob/ob* mice (Fig. 3 B and C). Administration of leptin reverted the changes in Drp1 expression and phosphorylation forms in *ob/ob* mice stimulating the Drp-1 inactive form (Figs. 3 A-C). In addition, we observed a reduction in Mfn2 and OPA-1 protein levels in *ob/ob*, which were restored after leptin treatment (Fig. 3 D and E). No changes were observed in wt C57BL/6 under any treatment.

**Figure 3.**
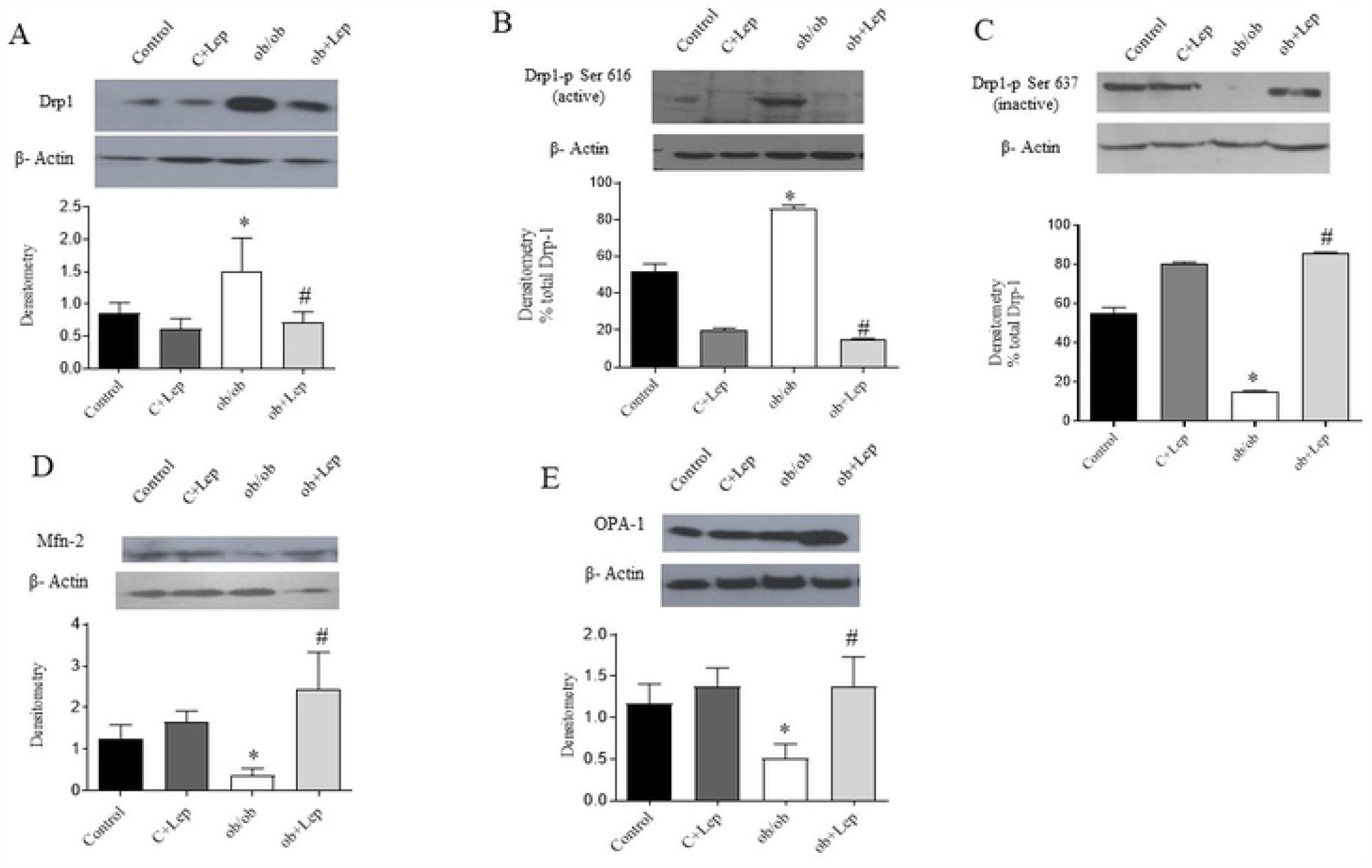
Effect of short-term leptin treatment on mitocondrial dynamics. (A) Representative immunoblot of proteins separated using SDS-PAGE from whole WAT lysate reveals the protein of (A) Drp1, (B) Drp-1 phosphorylated in Ser 616, bars reflect the Drp1-P protein quantified as a percentage of total DRP-1, (C) Drp-1 phosphorylated in Ser 637, bars reflect thep-Drp-1 protein quantified as a percentage of total Drp-1, and (D and E) the expression of fusion proteins Mfn2 and Opa-1. Data are normalized to ²-actin for protein. Bars reflect the densitometry in arbitrary units (A.U.) in absolute amount of protein. ^*^p < 0.05 denotes different from respective *wt C57BL/6*. # p < 0.05 denotes different from respective *ob/ob*, vs *ob/ob* + leptin treatment (1 mg/kg i.p). Results are mean ± SEM, n: 7 per group. One-way analysis of variance (ANOVA) and Bonferroni post hoc test.

### Leptin and mdvi-1 increase the expression of PGC-1*α* and AMPK-P as well the mitochondrial mass in WAT from *ob/ob* mice

After observing an overexpression of Drp-1 protein in *ob/ob* mice and to determine the effect of leptin or mdvi-1 treatment in mitochondrial biogenesis, we assessed PGC-1α gene expression. The mRNA level of PGC-1α was significantly lower in *ob/ob* mice compared to *wt C57BL/6*, leptin or mdvi-1 treatment (Fig 4 A). mtDNA content showed a reduction in *ob/ob* mice compared to *wt C57BL/6*. Inhibition of Drp-1 by mdvi-1 led to increased mtDNA content indistinguishable from leptin (Fig. 4 B). WAT obtained from *ob/ob* mice revealed that AMPK expression was devoid in its phosphorylated form (active) as compared to *wt*, while leptin or mdvi-1 treatment increased AMPK-P protein expression (Fig. 4 C and D).

**Figure 4.**
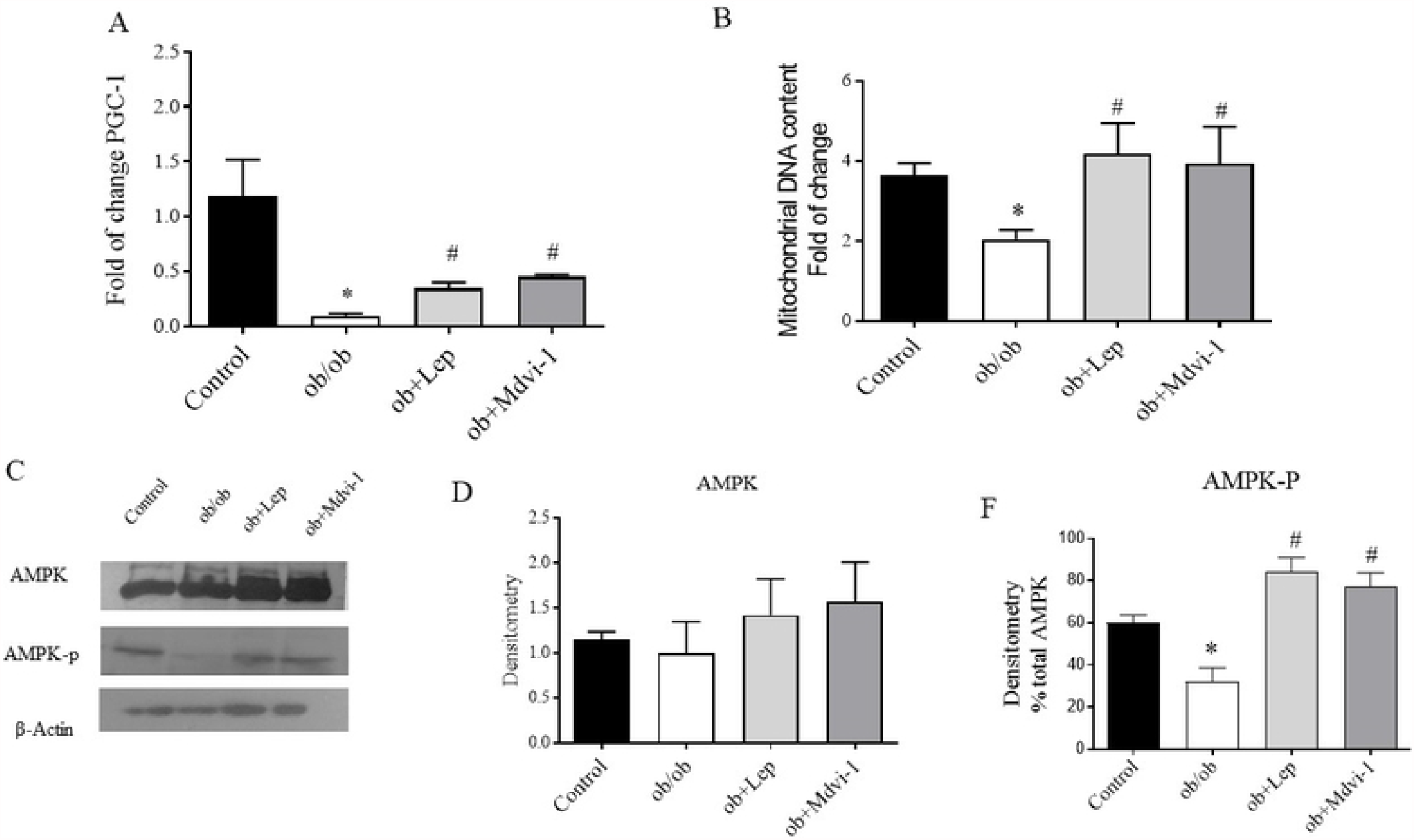
Effect of leptin and mdvi-1 on AMPK-P and PGC-1 *α* expression. (A) The mRNA levels of PGC-1 α were measured in TRIZOL-treated WAT extracts from mice treated with leptin or mdvi-1. Data are normalized to ² -actin for protein or GAPDH for mRNA. (B) Quantification of mtDNA and nuclear DNA content by qPCR in VAT in *wt C57BL/6, ob/ob* and *ob/ob* + leptin or mdvi-1 treatment, mtDNA/nDNA ratio was calculated. (C) Representative immunoblot of proteins separated using SDS-PAGE from whole WAT lysate reveals the expression of AMPK and AMPK-P in *wt, ob/ob, ob/ob* + leptin or mdvi-1. (D) Bars reflect the AMPK protein densitometry in arbitrary units (A.U.). (C) Bars reflect AMPK-P protein quantified as a percentage of total AMPK. (Results are mean ± SEM, n: 6-7 per group. ^*^p<.0.5 *wt C57BL/6* vs *ob/ob*, # p < 0.05 *ob/ob* vs *ob* + leptin or mdvi-1. One-way analysis of variance (ANOVA) and Bonferroni post hoc test.

### Drp1 inhibition increases the oxidative metabolism and mitochondrial function in WAT isolated adipocytes from *ob/ob* mice

To study the ^14^[C] palmitic acid and D-glucose ^14^[C] oxidation *in vitro*, isolated adipocytes from *wt C57BL/6* and *ob/ob* mice were incubated with leptin, mdvi-1 or siRNADrp-1. Basal palmitic acid oxidation in adipocytes from *ob/ob* was ∼20% of the *wt* (Fig. 5 A). Leptin, mdvi-1 and siRNADrp-1 significantly increased palmitic acid oxidation in *ob/ob* (3.8 pmol/min./mg.protein, 3.3 pmol/min./mg.protein, and 2.5 pmol/min./mg.protein, respectively) compared to *ob/ob* basal level (1.5 pmol/min./mg.protein) (Fig. 5 A). Likewise, leptin, mdvi-1 and siRNADrp-1 significantly increased glucose oxidation in *ob/ob* (20 nmol/min./mg.protein, 22 nmol/min./mg.protein, and 26 nmol/min./mg.protein, respectively) compared to *ob/ob* basal level (5.4 nmol/min./mg.protein) (Fig 5 B).

**Figure 5.**
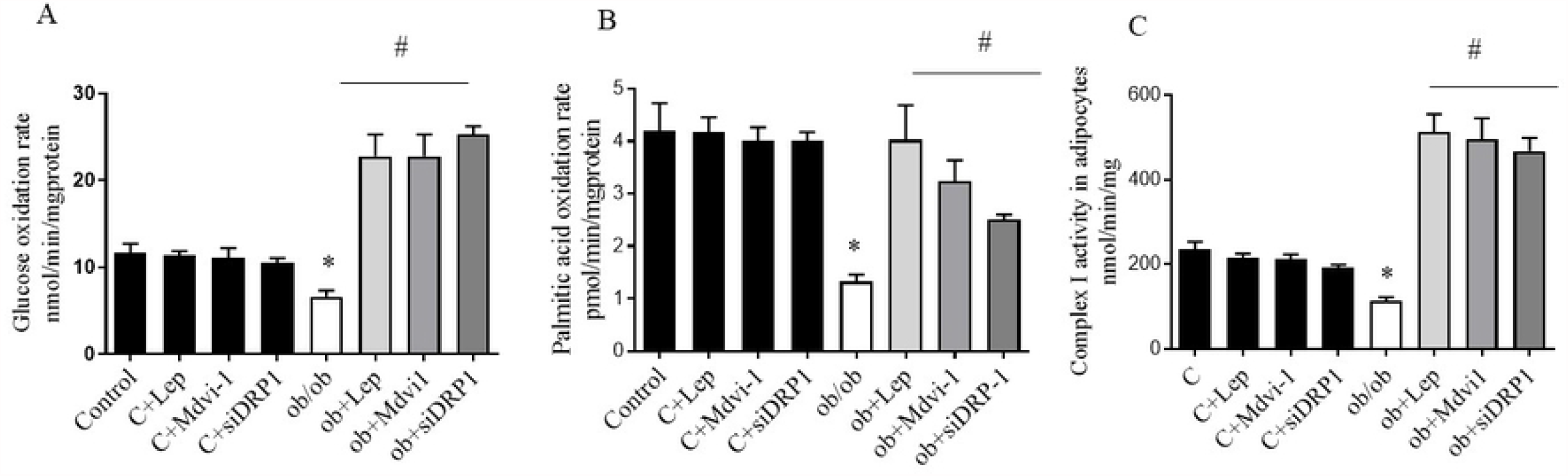
Effect of Drp-1 blocking on oxidative metabolism and mitochondrial function in isolated adipocytes from WAT. (A) Adipocytes from *wt* and *ob/ob* mice were isolated and incubated with leptin, mdvi-1 or siRNA DRP1 and fat metabolism was measured as fatty acid oxidation with 1μCi/ml ^14^[C]-palmitic acid oxidation to CO_2_ and H_2_O. Data resulted from pmolar/min./mg protein (Eli Wallac,;n= 5-6 per triplicate). (B) Glucose oxidation was measured with 1μCi/ml ^14^[C]-complete glucose oxidation to CO_2_ and H_2_O. Data resulted from nmolar/min./mg protein (Eli Wallac, n= 5-6 per triplicate). (C) Mitochondrial complex I activity was measured following the reduction of cytochrome c at 30 °C in the presence of NADH and KCN. The reaction rate was measured as the pseudo – first-order reaction constant (k’) and expressed as k’ /min. mg protein. test. Results are mean ± SEM, n: 5-6 per group. ^*^p < 0.05 *wt C57BL/6* vs *ob/ob*, # p < 0.05 *ob/ob* vs *ob/ob* + leptin or mdvi-1. One-way analysis of variance (ANOVA) and Bonferroni post hoc test.

The activity of the mitochondrial electron transport chain (ETC) complex I was evaluated in mitochondria. An impairment of mitochondrial function was observed in *ob/ob* (−50%), while leptin, mdvi-1 and siRNADrp-1 caused a 5-fold increase (Fig. 5 C). No changes were found in *wt C57BL/6* under any treatment.

### Effect of leptin and mdvi-1 in blood glucose level, body weight and food intake

Both leptin and mdvi-1 decreased the blood glucose concentration by 50% in *ob/ob* mice almost to the *wt* level. We did not observe any changes in body weight and food intake over the course of the treatments (Table 1).

**Table 1:** Effects of short-term leptin and mdvi-1 treatment on blood glucose levels, body weight and food intake. Blood glucose concentration (mg/dl) was measured in all studied groups. Body weight (g) and food intake (g/d) were determined in all animals during the study. Values are means ± SD of 8 mice per group ^*^p<.0.5 *wt C57BL/6* vs *ob/ob*, #p < 0.05 *ob/ob* vs *ob* + leptin or mdvi-1. One-way analysis of variance (ANOVA) and Bonferroni post hoc.

| Variable | C | Ob/ob | Ob + Lep | Ob + Mdvi-1 |
| --- | --- | --- | --- | --- |
| Blood glucose levels (mg/ml) | 138 ± 42 | 393 ± 36 * | 198 ± 22 ≠ | 178 ± 35 ≠ |
| Body Weight (g) | 25 ± 4 | 70.6 ± 6 * | 66 ± 4 * | 65 ± 7 * |
| Food Intake (g/d) | 2.4 ± 0.5 | 4.4 ± 0.4 * | 3.8 ± 0.3* | 3.5 ± 0.4* |

### Leptin and mdvi-1 stimulate the “browning” process in WAT from *ob/ob* mice

In order to study whether the leptin or mdvi-1 treatments in *ob/ob* mice stimulate the “browning” process in WAT, we measured the adipocyte area and vascular density and observed that leptin and mdvi-1 treatment reduced the adipocyte area and increased the vascular density (Fig. 6 A and B). In addition, we determined that mRNA UCP-1 expression was increased by mdvi-1 more than leptin in *ob/ob* WAT (Fig. 6 C).

**Figure 6.**
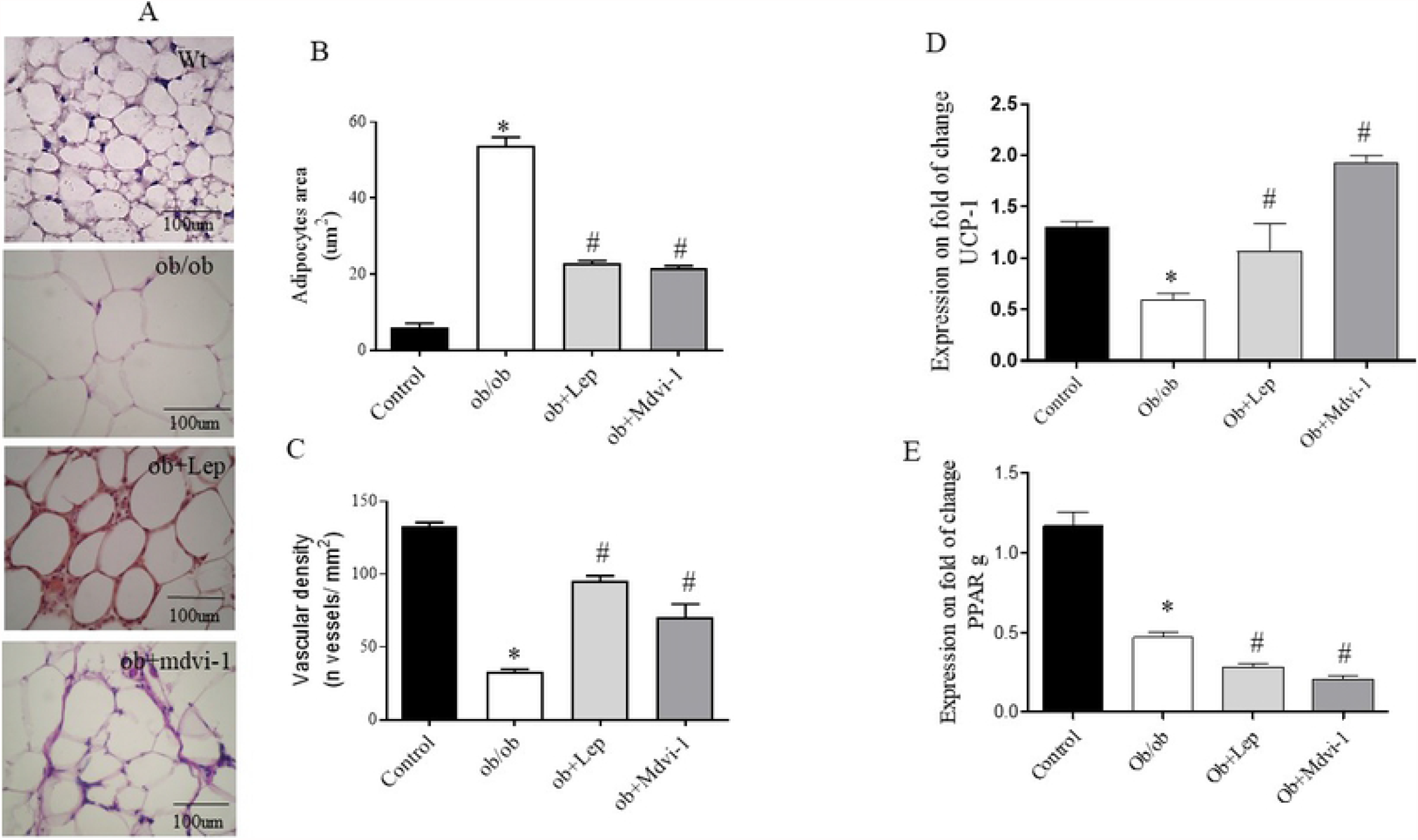
Effect of leptin and mdvi-1 on browning process on WAT. (A) Representative micro-sections of WAT were stained with hematoxylin and eosin reagent and periodic acid-schiff (PAS) stain for studying the adipocytes area (B) and vascular density adipocyte (C) in all studied groups. (D) The mRNA levels of UCP-1and (E) PPARγ were measured in Trizol-treated WAT extracts from all the studied groups; GADPH mRNA was used as standard (n 5-6). Results are mean ± SEM, n: 6-7 per group. ^*^ p < 0.05 *wt C57BL/6* vs *ob/ob*, # p < 0.05 *ob/ob* vs *ob/ob* + leptin or mdvi-1. One-way analysis of variance (ANOVA) and Bonferroni post hoc test.

## DISCUSSION

This study focuses on mitochondrial dynamics and biogenesis as both processes are involved in WAT in obesity and diabetes and their modulation by leptin and mdivi-1.

Mitochondrial fragmentation and mitophagy prevail in *ob/ob* mice WAT; however, both mdivi-1 and leptin treatment increase the percentage of tubular mitochondria (fusion) and decrease the amount of fragmented mitochondria (fission), disclosing that the defects observed in mitochondrial morphology are linked to fission activation. Mitophagy plays an important part in removing damaged mitochondrial structure, which is facilitated by prior fission (32). We found a high number of images compatible with mitophagy in WAT from *ob/ob* mice. During autophagosome formation, cytosolic LC3-I is conjugated to phosphatidylethanolamine with the resulting LC3-II localized to autophagosomes (33). According to that, we observed an increase in LC3 II protein expression. This abnormality in fusion-fission-mitophagy ratio improves with leptin and mdvi-1. To confirm matching changes of mitochondrial morphology in cells, we studied the effect of leptin or mdvi-1 in isolated adipocytes from treated mice, and we observed an increment of longer mitochondria, as a result of increased fusion and decreased fission. The stronger pro-fusion and anti-fission effect of leptin and mdvi-1 could explain the proportional increase of complex shaped mitochondria as observed in the “branched mitochondria” group.

Upon observing the presence of mitochondrial fragmentation and mitophagy in WAT from *ob/ob* mice and its modulation by leptin and mdvi-1, we studied their effect on mitochondrial dynamics. We observed an increase in the expression of mitochondrial fission-protein level in *ob/ob* respect to *wt* WAT (total Drp-1 and Drp1-pSer 616 active isoform), while concomitantly reducing the expression of proteins responsible for mitochondrial fusion (Mfn2 and OPA-1).

The mechanism of controlling mitochondrial biogenesis, promoting mitochondrial DNA replication and integrity, and increasing insulin sensitivity is associated with PGC-1α (34,35). Our data show a decreased mRNA PGC-1α expression and mtDNA/nDNA ratio in *ob/ob*, while leptin and mdivi-1 induce mitochondrial biogenesis increasing PGC-1α levels and mitochondrial mass. In agreement with our data, evidence supports alterations in mitochondrial dynamics and reduced expression of PGC-1α and Mfn2 resulting in the loss of mitochondrial mass and structure abnormalities in obesity and type 2 diabetes in humans, rats, leptin-deficient *ob/ob* and high-fat-diet-fed dietary mice with hyperleptinemic state (36-38).

The short-term leptin hormone replacement in hypoleptinemic state is not only sufficient to normalize the levels of biogenesis and dynamics proteins but also inactivates Drp-1 by Ser 637 phosphorylation.

The accumulation of three major nutrients –glucose, fat acids and aminoacids– in obesity and diabetes with glucose levels suppresses AMPK through mechanisms that do not affect the AMP/ATP ratio and contributes to insulin resistance, oxidative stress and mitochondria dysfunction to impede substrate oxidation and to promote fat accumulation and obesity (39). It has also been reported that AMPK activation indirectly phosphorylates Drp-1 at Ser 637, and this phosphorylation has been linked to the inhibition of Drp-1 and a decrease in mitochondrial fission. In addition, AMPK and aerobic exercise downregulate the phosphorylation level of Drp1 at Ser 616 which is upregulated by ROS. Likewise, AMPK plays a role in mitochondrial homeostasis phosphorylating PGC-1α on specific serine and threonine residues leading to an increased mitochondrial gene expression. AMPK is activated by leptin in the peripheral tissues increasing fatty acid oxidation and glucose uptake. Leptin produces Drp-1 inhibition modulating mitochondrial fission and PGC-1α stimulation favoring mitochondrial biogenesis; both are protective effects mediated by AMPK (40,41). In our study, leptin and mdvi-1 induce AMPK-P supporting the idea that AMPK can work in the context of different cellular signals that are distinguished from energetic demand, to promote a variety of mitochondria function.

Elongated and branched mitochondria have been previously associated with improved oxidative phosphorylation and several bioenergetic advantages, while round mitochondria have been associated with a predominance of oxidative stress and dysfunction (42). In the present study, we detected decreased complex I activity and oxidative metabolism in *ob/ob* adipocytes, counteracted by leptin, mdvi-1 and siRNAs Drp-1, which increased complex I activity followed by an increment of glucose and palmitic acid oxidation, depicting that mitochondrial function is closely related to Drp-1 inhibition. In accordance with the metabolic and mitochondrial effects, leptin and mdvi-1 treatments produce a significant decrease in blood glucose levels in *ob/ob* mice which typically exhibit high levels of blood glucose concentration. Normalization of blood glucose levels were not accompanied by a decrease in body weight and food intake, suggesting that blood glucose concentration was related to increased glucose oxidative metabolism secondary to changes in mitochondrial homeostasis.

“Browning” or “beiging” WAT process contributes to high metabolic rates and a high level of mitochondrial content in these cells. UCP-1-mediated thermogenesis is a hallmark of brown/beige adipocytes development (43). According to numerous cold exposure studies in mice, subcutaneous adipose tissue depots are more susceptible to browning than the metabolically unfavorable visceral fat. However, it has been reported that in obese subjects, omental (visceral) fat expressed higher transcript levels of browning markers and genes involved in mitochondrogenesis compared to abdominal subcutaneous fat (44). We studied whether epididymal white adipose tissue would be capable of undergoing the browning process. The adipocytes termed “beige” or “brite” (brown in white) show an intermediate phenotype between white and brown adipocytes, exhibiting multilocular morphology, high levels of UCP-1 expression and higher vascularization (45), as seen in epididymal white adipose tissue in *ob/ob* with leptin and mdivi-1 treatment. Hypoleptinemic and hyperleptinemic state and the loss of AMPK-P in WAT in obesity and diabetes are likely a driving force for re-whitening (inactivation) of beige adipocytes, exacerbating insulin resistance through impaired brown and beige fat function (46).

Leptin production by adipocytes is critical for energy homeostasis activating AMPK in peripheral tissues and is the promoter of BAT thermogenesis in both UCP-1-mediated and independent thermogenesis mechanisms (47). Leptin acute and chronic effects have been studied in different models and pathological situations. While short-term leptin exposure produces lipolytic effects, a lipogenic effect has been observed with leptin chronic stimulus in mature adipocytes (48). Some studies reported that leptin directly stimulated lipolysis in adipocytes isolated from fasted wild type or *ob/ob* mice (49). There is also evidence that lipolysis inhibition in adipose tissue under longer periods of leptin exposure reduced endothelial cell expression of PPARγ, while chronic leptin treatment restored PPARγ expression in mice (50). We hypothesize that in mature adipocytes short-term leptin and mdvi-1 treatment directly induces lipolysis inhibiting PPARγ expression. In summary, mitochondrial dysfunction has been related with obesity and type 2 diabetes, considered a modern epidemic. *Ob/ob* mice is a model of leptin deficiency with hyperinsulinemia, reduced levels of AMPK-P and high levels of mitochondrial fission and mitophagy that induce mitochondrial dysfunction and increase oxidative and nitrosative stress in WAT. The inhibition of mitochondrial fission by leptin and mdvi-1 stimulates mitochondrial biogenesis, fusion and oxidative capacity. Modulation of mitochondrial network through Drp-1, a critical mediator of mitochondrial fission, would have an important role in browning white adipose tissue into beige adipocytes. We suggest that Drp-1 blocking can be now proposed as a novel therapeutic target for obesity and diabetes. Based on that, it seems reasonable to predict that future new therapies targeting mitochondrial fission could be tested as potential alternatives.

## Abbreviations Used

AMPK: AMP-dependent kinase
BAT: brown adipose tissue
DAF-FM: amino-5-methylamino-2¢,7¢-difluorofluorescein diacetate
DRP-1: dynamin-related protein-1
FFA: free fatty acid
LC3-II: Microtubule-associated protein 1A/1B-light chain 3
Mdivi-1: selective inhibitor of Drp1
Mfn2: mitofusin proteins
NO: nitric oxide
NRF1: nuclear respiratory factor
Ob/ob: leptin-deficient mice
OPA-1: optic atrophy-1
OXPHOS: oxidative phosphorylation system
PE: phosphatidylethanolamine
PGC-1α: proliferator-activated receptor gamma coactivator-1 alpha
RNS: reactive nitrosative species
ROS: reactive oxygen species
siRNA: small interference RNA
UCP-1: uncoupler protein 1
WAT: white adipose tissue

## Acknowledgments

The authors thank Margarita López, Fabiana Confente and Mariana López Ravasio for electron microphotographs, Inés Rebagliatta and Silvia Holod for technical support; and Natalia Riobó for revising of the manuscript.

## Author Contribution

P.F., M.C.C., and J.J.P. designed the study; P.F. performed experiments; H.P. performed the RNA extraction, Quantitative Real-Time PCR, and mtDNA content experiment; G.B. performed fluorescence microscope experiment, V.M. and G.B. performed optic microscope and area, density and vascular white adipose tissue study, C.M. performed technical assistant, C.M. performed histological study, P.F., and M.C.C. collected and analyzed data; P.F., H.P., J.P., C.P., J.J.P. and M.C.C. wrote the manuscript.

## SUPPLEMENTAL MATERIAL

**Supplemental Table I:**
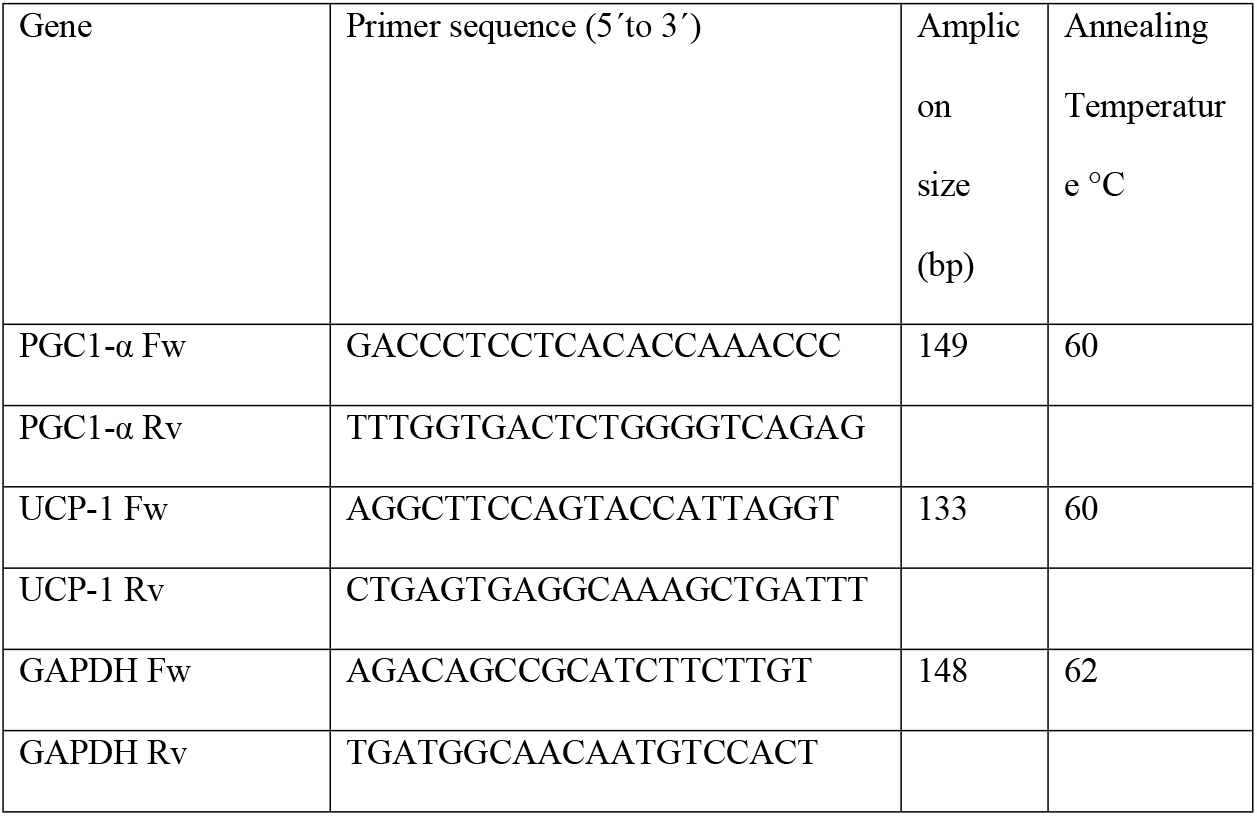
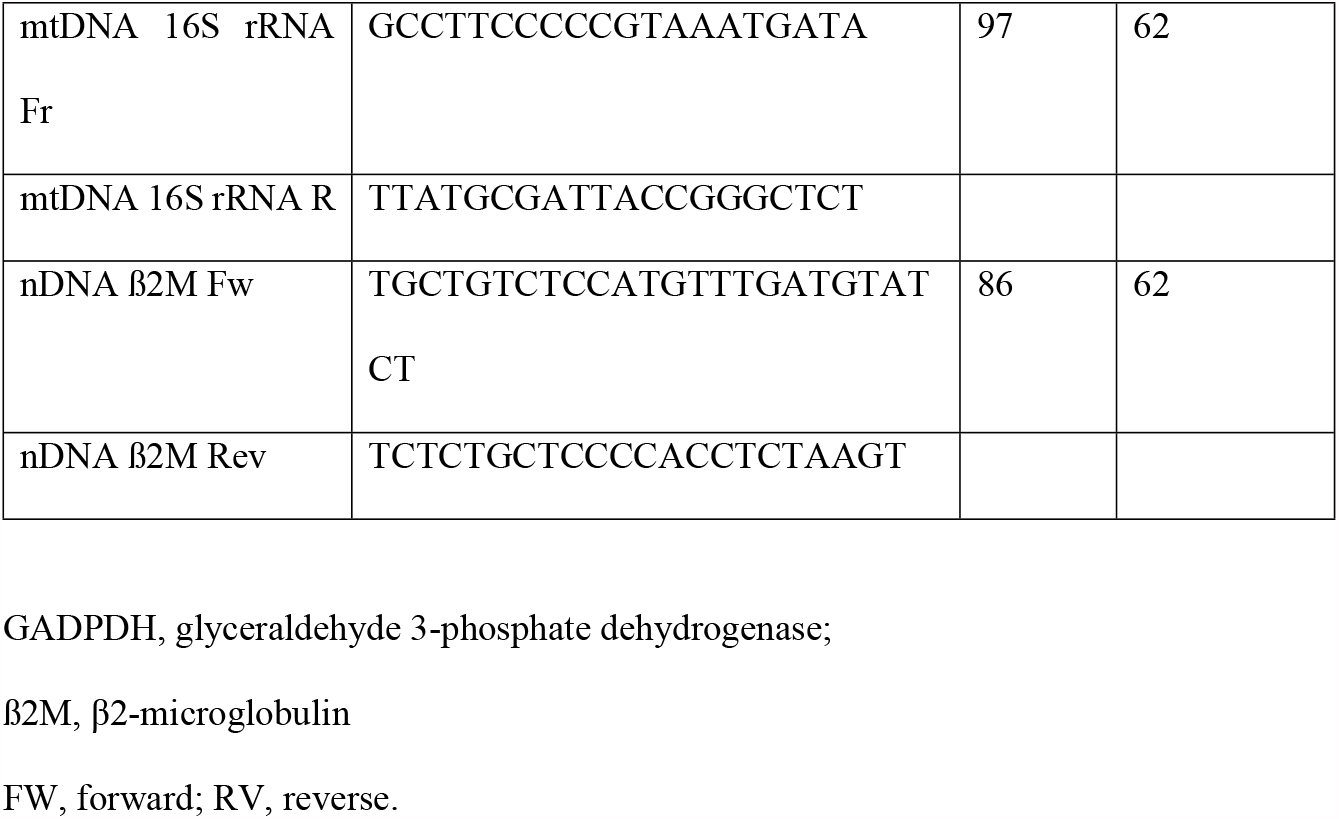
Mice gene-specific oligonucleotide sequences used in quantitative real-time polymerase chain reaction.

**Supplemental Figure 1: Assessment of the fraction of mitochondria shape-classes by supervised classification**

A Cell Profiler pipeline was created for the segmentation and measurement of morphometric parameters of individual mitochondria. Several images were processed with this pipeline for each treatment and morphometric parameters were exported and processed with Cell Profiler Analyst (Broad Institute, USA). A classification algorithm was used to implement a machine learning approach to classify mitochondria intro three groups: round, elongated and branched. The training procedure was repeated until global classification accuracy achieved 90%, with 100% classification accuracy of the round mitochondria subgroup (further details a references in methods section).

(A) An example of a supervised classification of a single image used to verify the ability to discriminate among morphological classes. Different mitochondrial colours mean mitochondrial morphology: blue and light blue: a branched, yellow: rounded, orange and red:elongated.

(B) A sample of the training set used for machine learning and the confusion matrix obtained after training. The matrix was used to confirm the accuracy of the scoring of the whole set of mitochondria.

**Supplemental Figure 2: Expression of Drp-1 after siRNA transfection and dose-response of leptin on p-STAT3 expression in adipocytes**

(A) Representative immunoblot of proteins separated using SDS-PAGE from whole WAT lysate reveals the expression of Drp-1 after transfection of *ob/ob* adipocytes with empty-vector and siRNA Drp-1 (50nM), respectively.

(B) Dose-dependent effect of leptin on the expression of p-STAT3 (Ser^727^) in *ob/ob* adipose tissue.

